# Heterogeneity of social cognition in temporo-parietal junction: Overlapping yet distinct representation between visual perspective-taking and theory of mind

**DOI:** 10.1101/2022.01.04.474884

**Authors:** Kenji Ogawa, Yuiko Matsuyama

## Abstract

Visual perspective taking (VPT), particularly level 2 VPT (VPT2), which allows an individual to understand that the same object can be seen differently by others, is related to the theory of mind (ToM), because both functions require a decoupled representation from oneself. Although previous neuroimaging studies have shown that VPT and ToM activate the temporo-parietal junction (TPJ), it is unclear whether common neural substrates are involved in VPT and ToM. To clarify this point, the present study directly compared the TPJ activation patterns of individual participants performing VPT2 and ToM tasks using functional magnetic resonance imaging and within-subjects design. VPT2-induced activations were compared with activations observed during a mental rotation task as a control task, whereas ToM-related activities were identified with a standard ToM localizer using false-belief stories. A whole-brain analysis revealed that VPT2 and ToM activated overlapping areas in the posterior part of the TPJ. By comparing the activations induced by VPT2 and ToM in individual participants, we found that the peak voxels induced by ToM were located significantly more anteriorly and dorsally within the bilateral TPJ than those measured during the VPT2 task. We further confirmed that these activity areas were spatially distinct from the nearby extrastriate body area (EBA), visual motion area (MT+), and the posterior superior temporal sulcus (pSTS) using independent localizer scans. Our findings revealed that VPT2 and ToM have distinct representations, albeit partially overlapping, indicating the functional heterogeneity of social cognition within the TPJ.

**Significance Statement:** The temporo-parietal junction (TPJ) is consistently activated by social cognitive tasks such as visual perspective taking (VPT) and theory of mind (ToM) tasks. The present study investigated whether VPT and ToM have the same neural substrates within the TPJ using functional magnetic resonance imaging (fMRI) with a within-subjects design. While VPT and ToM tasks activated overlapping areas in the posterior part of the TPJ, the individual peak voxels induced by ToM were located significantly more anteriorly and dorsally compared with those observed during the VPT task. Moreover, they were spatially distinct from the nearby functional modules, such as the extrastriate body area, visual motion area, and posterior superior temporal sulcus. Our findings reveal the heterogeneity of social cognition representation within the TPJ.

## Introduction

Visual perspective taking (VPT) is the ability to understand how the world looks to others and has two different levels. Level 1 VPT (VPT1), also called perspective tracking (Gunia et al., 2021), is the ability to determine if another person can see an object, whereas level 2 VPT (VPT2) is the ability to understand that others have different views of an object than one’s own (Flavell, 1977). Previous studies have indicated that VPT1 and VPT2 are distinct cognitive processes. Developmental studies have shown that VPT1 is acquired between 18 and 24 months of age, whereas VPT2 appears in 4- or 5-year-old children (Flavell et al., 1981; Gzesh and Surber, 1985; Pearson et al., 2013). Behavioral investigations also support that VPT1 and VPT2 have different properties. In VPT2, the response times increase with the increase in the angular distance between the participant and the agent, indicating that the participant mentally transformed their own position to the other’s perspective (Michelon and Zacks, 2006). Studies using mental simulation of body movements also proposed that VPT2 is an embodied process (Kessler and Rutherford, 2010; Hirai et al., 2018).

VPT2 is related to the theory of mind (ToM) or mentalizing, because both functions require a decoupled representation from one’s own perspective or belief. Moreover, people with autism spectrum disorder (ASD) present impaired ToM and VPT2, whereas the VPT1 ability remains intact (Hamilton et al., 2009; Pearson et al., 2013). Hamilton et al. (2009) compared the performance of children with ASD with those of typically developed children in terms of the VPT2 and in mental rotation (MR) task, both of which require comparable mental spatial processing. They showed that VPT2, but not MR, performances were significantly impaired in the children with ASD compared with those of typically developed children. They also showed that VPT2 performances were significantly associated with the ToM ability (Hamilton et al., 2009). Their findings indicate that VPT2 and ToM have common cognitive substrates.

The temporo-parietal junction (TPJ) has been involved in both VPT and ToM. Lesions in the TPJ induce impairments of the ToM, particularly of the ability to understand other’s false beliefs (Apperly et al., 2004; Samson et al., 2004) and of VPT, especially the ability of taking another person’s perspective (Samson et al., 2005). In addition, transcranial direct stimulation of the TPJ improves VPT performances in normal subjects (Santiesteban et al., 2015; Martin et al., 2020). Many neuroimaging studies have consistently shown TPJ activations during the processes of VPT (Aichhorn et al., 2006; Corradi-Dell’Acqua et al., 2008; Schurz et al., 2015; Agarwal et al., 2017) and ToM (Saxe and Kanwisher, 2003; Otsuka et al., 2009; Dodell-Feder et al., 2011; Schurz et al., 2014; Kanske et al., 2015). Some meta-analyses have also shown that VPT and ToM cause TPJ activation (Schurz et al., 2013; Gunia et al., 2021). However, another study found no overlap activation between VTP and the different types of ToM tasks (Arora et al., 2017). Although VPT and ToM activate regions within the TPJ, they might not share the same neural substrates, considering that activations within the same region of interest do not necessarily mean common neural substrates (Hong et al., 2019). Since the TPJ is heterogeneous and composed of various subregions (Decety and Lamm, 2007; Mars et al., 2012; Bzdok et al., 2013), it is important to investigate the common neural substrates between VPT and ToM by testing the regions activated within individual participants.

The present study aimed to investigate whether VPT and ToM share common neural substrates in the TPJ using functional magnetic resonance imaging (fMRI) and within-subjects design. Participants performed VPT and ToM tasks. The regions activated by VPT were compared with those activated by an MR task as the control task. The activities triggered by ToM were identified with a standard ToM localizer. The activations were then compared within participants to identify potential shared neural substrates for VPT and ToM in the TPJ.

## Materials and Methods

### Participants

Participants included 31 healthy volunteers (21 males and 10 females; 20–27 years of age). The number of participants was determined to detect reliable activations based on previous fMRI experiments on VPT (Aichhorn et al., 2006; Schurz et al., 2015) or ToM (Saxe and Kanwisher, 2003; Young et al., 2010b). All participants, with the exception of one, were right-handed, as assessed by a modified version of the Edinburgh Handedness Inventory (Oldfield, 1971) for Japanese participants (Hatta and Nakatsuka, 1975). Written informed consents were obtained from all participants in accordance with the Declaration of Helsinki. The experimental protocol was approved by the local ethics committee.

### Task procedures

Each participant underwent a VPT and a ToM task. For the VPT task, we employed a computerized version of the paradigm by Hamilton et al. (Hamilton et al., 2009) for VPT and MR tasks for children. Although their original study used a real meaningful object (small toy), we used pseudo-randomly combined three-dimensional cubes typically used for MR (Shepard and Metzler, 1971) to increase the task difficulty for adults. First, one object and a pot, which were placed on a tray whose front side was colored red, were displayed on the left and right sides of the screen, respectively, for 3 s. In addition, in the VPT condition, an image of a person (avatar) standing either on the front, back, left, or right side of the pot was displayed. The participants were asked to view the object from the perspective of the avatar. In the MR condition, one of the four sides of the tray was colored red. The participants were asked to imagine the tray rotating in the direction indicated by the red side of the tray as well as the rotation of the object. In both VPT and MR conditions, four different views of the same object were presented and the participants had 4 s to select the correct response (Fig. 1). Between the trials, a fixation cross was presented for 8 s. The VPT and MR trials were pseudo-randomly presented in the same run, with 20 trials per condition. One run lasted for approximately 10 min.

**Fig. 1:**
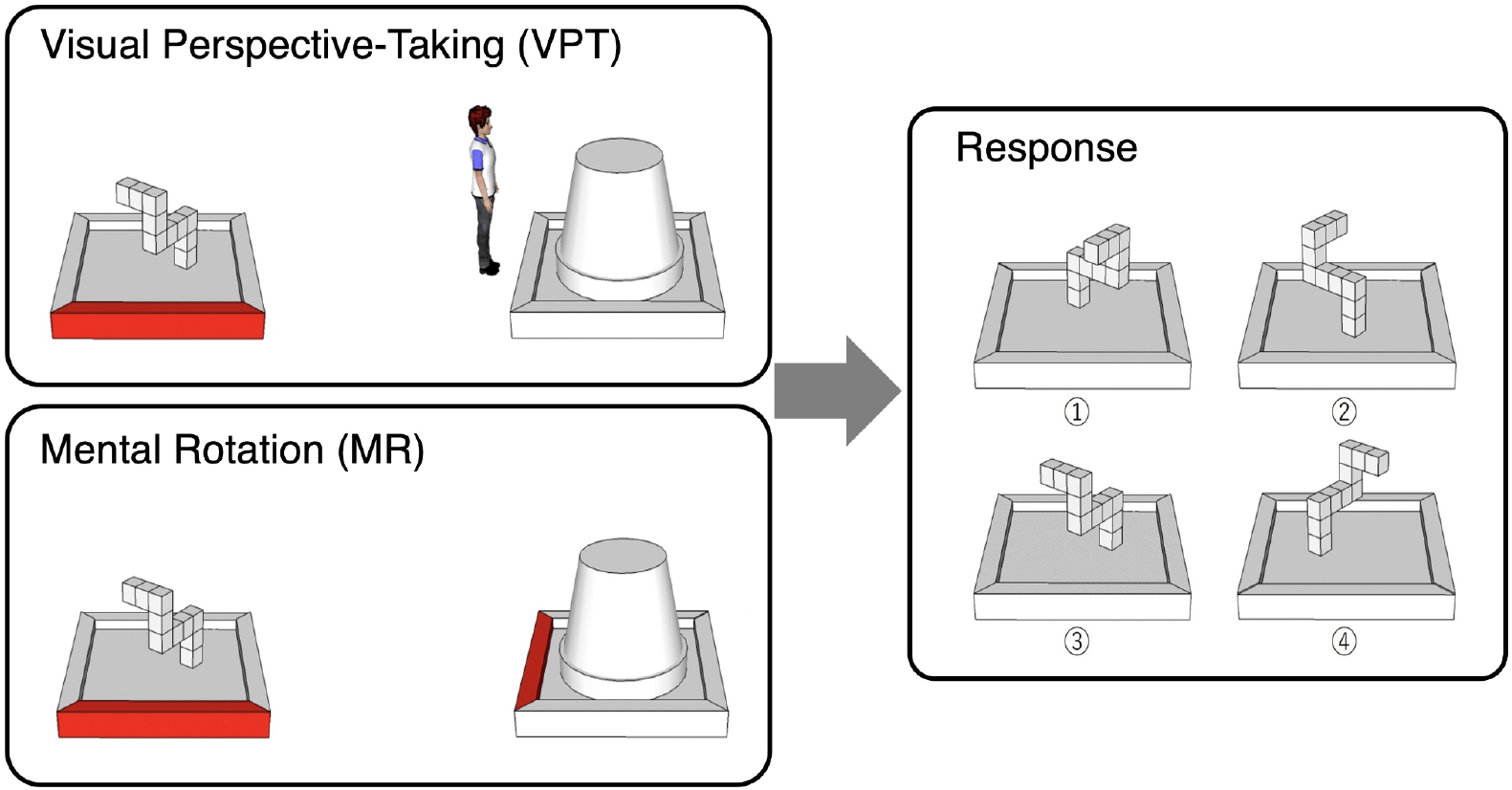
Stimuli used for the VPT and MR tasks. First, a 3D object and a pot, which were placed on a tray with the front side colored red, were displayed for 3 s. In addition, an image of a person (avatar) standing either on the front, back, left, or right side of the pot was displayed in the VPT condition. The participants were asked to imagine the view of this object from the perspective of this person. In the MR condition, one of the four sides of the tray was colored in red. The participants were asked to image the rotation of the tray in the direction indicated by the red side of the tray as well as the rotated view of the object. In both VPT and MR conditions, four different views of the same object were then presented, and the participants had 4 s to select the correct response.

For the ToM task, we used a modified version of standard ToM localizer (Dodell-Feder et al., 2011) for Japanese participants (Ogawa et al., 2017). The participants were presented with stories describing false beliefs in the ToM task or false pictures or maps in the control task. Both types of stories were pseudo-randomly ordered within a run, with seven stories per condition. Each story was displayed as written sentences for 10 s. After the story, a true or false question asking whether the situation was real or was a false representation was presented for 4 s. Between the trials, a fixation cross was displayed for 7 s. One run lasted for approximately 5 min.

VPT and ToM tasks were conducted in separate runs and each participant underwent a total of three runs, two VPT runs and one ToM run. The order of the VPT and ToM runs was counterbalanced across participants. The participants’ response (pressing a button) and reaction time were recorded using an MRI-compatible response pad (Current Design, Philadelphia, USA). The stimuli were designed using SketchUp Make 2017 (Trimble Inc., California, USA), and the programs were implemented with PsychoPy 3 (https://www.psychopy.org/).

### Magnetic resonance imaging (MRI) acquisition

All scans were performed on a Siemens (Erlangen, Germany) 3-Tesla Prisma scanner with a 64-channel head coil at Hokkaido University. T2*-weighted echo planar imaging (EPI) was used to acquire a total of 304 scans for the VPT runs and 151 scans for the ToM runs, with a gradient echo EPI sequence. The first three scans within each run were used for T1 equilibration and were discarded. The scanning parameters were repetition time (TR): 2,000 ms, echo time (TE): 30 ms, flip angle (FA): 90°, field of view (FOV): 192 × 192 mm, matrix: 94 × 94; 36 axial slices, and slice thickness: 3.0 mm with a 0.75 mm gap. T1-weighted anatomical imaging with a magnetization-prepared rapid gradient-echo sequence was performed using the following parameters: TR: 2,300 ms, TE: 2.32 ms, FA: 8°, FOV: 256 × 256 mm, matrix: 256 × 256, 192 axial slices, and slice thickness: 1 mm without a gap.

### Processing of fMRI data

Image preprocessing was performed using the SPM12 software (Wellcome Department of Cognitive Neurology, http://www.fil.ion.ucl.ac.uk/spm) in MATLAB R2021a (The MathWorks, Inc). All functional images were initially realigned to adjust for motion-related artifacts. Volume-based realignment was performed by co-registering images using rigid-body transformation to minimize the squared differences between volumes. The realigned images were then spatially normalized using the Montreal Neurological Institute (MNI) template based on the affine and non-linear registration of coregistered T1-weighted anatomical images (normalization procedure of SPM). They were resampled into 3-mm-cube voxels using sinc interpolation. Images underwent spatial smoothing using a Gaussian kernel of 6 × 6 × 6-mm full width at half-maximum.

Using the general linear model, the task blocks of each session were modeled as box-car regressors that were convolved with a canonical hemodynamic response function. For both VPT and ToM runs, the box-car covered the presentation period of the first stimuli (4 s for VPT and 10 s for ToM). The six realignment parameters were also included in the design matrix as covariates. Low-frequency noise was removed using a high-pass filter with a cut-off period of 128 s, and serial correlations among scans were estimated with an autoregressive model implemented in SPM12.

### fMRI analysis

We used the conventional mass-univariate analysis of individual voxels to identify the areas activated by finger tapping. For VPT runs, we analyzed areas that were significantly activated during the VPT task compared with those activated during MR trials (VPT > MR). For ToM runs, the regions that were significantly activated during the ToM task were compared with those activated in control trials (ToM > control). Contrast images of each participant, generated using a fixed-effects model, were analyzed using a random effects model of one-sample t-test. The activation was reported with a threshold of p < .05 corrected for multiple comparisons of family-wise error (FWE) at the cluster-level and with p < .001 uncorrected at the voxel level. In addition, to compare their activation patterns, individual voxel coordinates of peak activation were selected within a spherical region of interest (ROI) with a radius of 20 mm centered on the reported coordinates of the bilateral TPJ (right TPJ: −49, −61, 27; left TPJ: 54, −55, 20) based on a recent meta-analysis of neuroimaging studies for social cognition (Alcalá-López et al., 2018). The same ROI mask was applied for VPT and ToM analysis to ensure unbiased selection of the coordinates.

### Functional localizer scans

We additionally conducted standard functional localizer scans to identify the following three regions in each participant: the extrastriate body area (EBA) (Downing et al., 2001; Spiridon et al., 2006), visual motion area (MT+) (Tootell et al., 1995; Wilms et al., 2005), and the posterior superior temporal sulcus (pSTS) (Grossman et al., 2000; Peuskens et al., 2005). These three localizer runs were independently scanned for the 12 subjects (9 males and 3 females; 20–24 years of age) who participated in the main experiment. Each run lasted for approximately 7 min.

In the EBA localizer, one block consisted of 20 images of headless human body parts in different postures, which were alternated with 20 images of chairs as controls. Each image was presented for 300 ms, followed by a black screen for 450 ms. A fixation cross was intercalated at the end of each block for 12 s as baseline. The same images were presented twice in succession, twice during each block. The participants were asked to press a button with the right index finger when they detected the immediate repetitions. In the MT+ localizer, the viewing block consisted of presenting moving or static dots for 15 s alternately with a rest period of fixation display for 12 s. A total of 16 blocks, which consisted of 8 moving blocks and 8 static blocks, were executed. In the pSTS localizer, a point-light biological motion was displayed for 15 s. In the control condition, scrambled displays with the same motion vectors as those of the biological motion but with randomized initial starting positions were presented for 15 s. A total of 16 blocks were performed, which consisted of 8 biological motion blocks and 8 control blocks. The participants passively viewed the stimuli during the EBA and pSTS localizer scans. The threshold of the activation peak was set at p < .05 uncorrected for the multiple comparisons and selected within a spherical mask with a 20-mm radius centered on the previously published coordinates as follows: EBA coordinates were based on the mean coordinates of nine neuroimaging paper (Okamoto et al., 2017), whereas MT+ and pSTS were selected from the meta-analysis of social cognition (Alcalá-López et al., 2018).

## Results

### Behavioral analysis

We analyzed the task accuracy and the reaction time (RT) (Fig. 2). The mean (SD) accuracies in the VPT and MR tasks were 87.3% (14.5%) and 70.6% (20.0%), respectively. Results of the paired t-test showed that the accuracy in the VPT task was significantly higher than that in the MT task, t(30) = 5.07, p < .001, Cohen’s d = 0.93. The mean (SD) RTs in the VPT and MR tasks were 2.63 (0.59) and 2.71 (0.56) s, respectively. The VPT RTs were significantly shorter than those observed in the MR task, t(30) = 2.20, p < .05, Cohen’s d = 0.40.

**Fig. 2:**
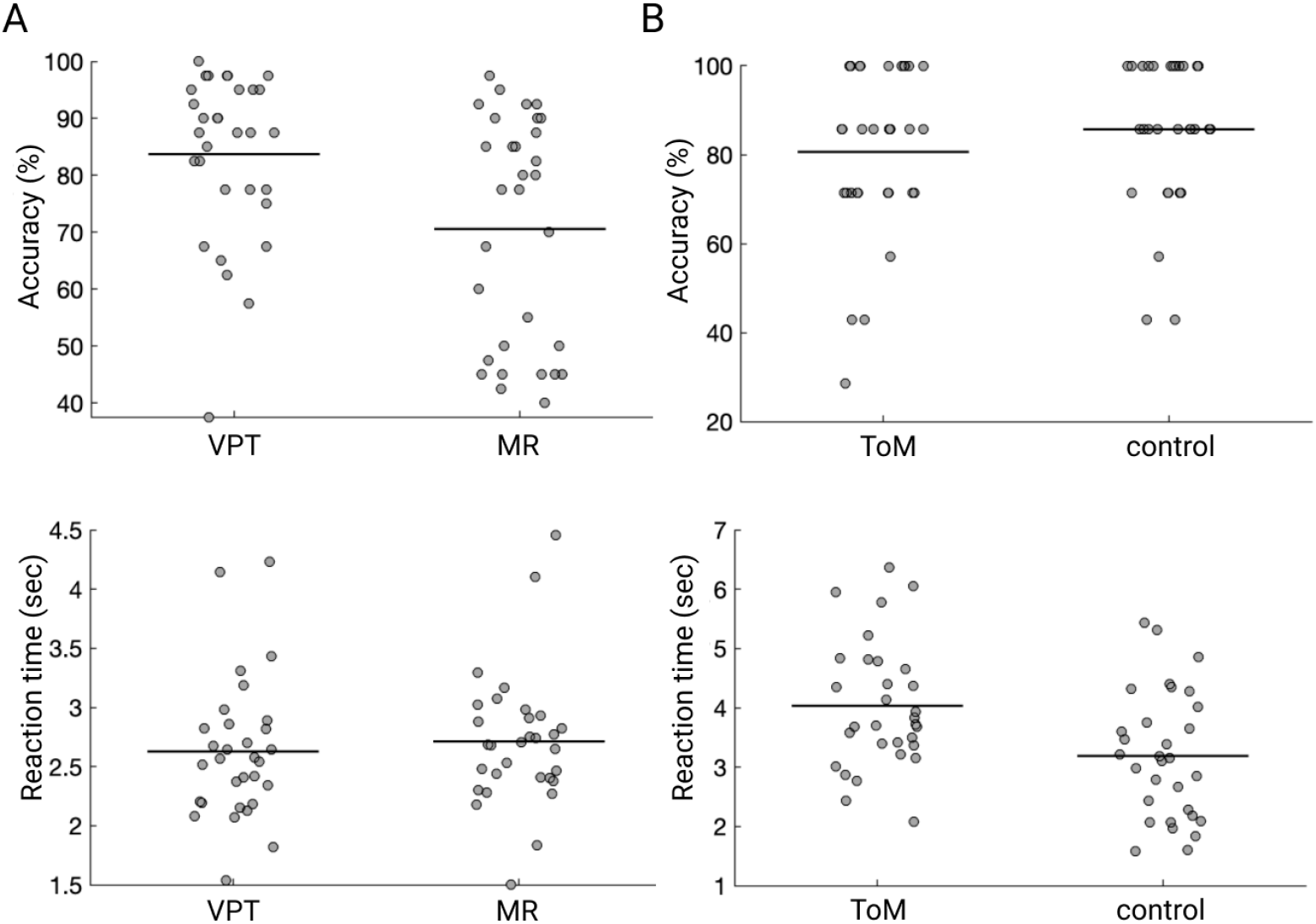
Behavioral results for each run. (A) Task accuracy and reaction time (RT) in the VPT and MR conditions. (B) Task accuracy and RT in the ToM and control conditions. Each dot represents the data of an individual subject. The horizontal bar indicates the mean value. The individual data are available at https://doi.org/10.6084/m9.figshare.17149094

The mean (SD) accuracy in the ToM localizer task and control condition were 80.6% (19.0%) and 85.7% (16.5%), respectively, without significant difference between both tasks, t (30) = 1.61, p = 0.12, Cohen’s d = 0.29. The mean (SD) RTs in the ToM localizer task and control condition were 4.03 (1.07) and 3.19 (1.07) seconds, respectively. The RTs in the ToM localizer task were significantly slower than those in the control condition, t (30) = 5.08, p < .001, Cohen’s d = 0.93.

### Analysis of fMRI data

A whole-brain analysis was conducted to identify the activations specific to the VPT and ToM conditions. We found notably higher activations in the bilateral TPJ during VPT tasks than in those observed in MR trials (VPT > MR). The regions preferentially activated during ToM compared with those activated in control condition were mostly in the bilateral TPJ extending into the anterior temporal cortex (ToM > control). Activations of the medial prefrontal cortex (mPFC), the precuneus, the early visual cortex, and the cerebellum were also detected (Fig. 3A, B, and Table 1). We then overlapped the voxels activated during the VPT and ToM tasks and found a relatively small overlap (blue voxels in Fig. 3C, number of voxels = 69) compared with the original cluster size activated by VPT (Fig. 3A in red, number of voxels = 3,124) and ToM (Fig. 3B in green, number of voxels = 130).

**Fig. 3:**
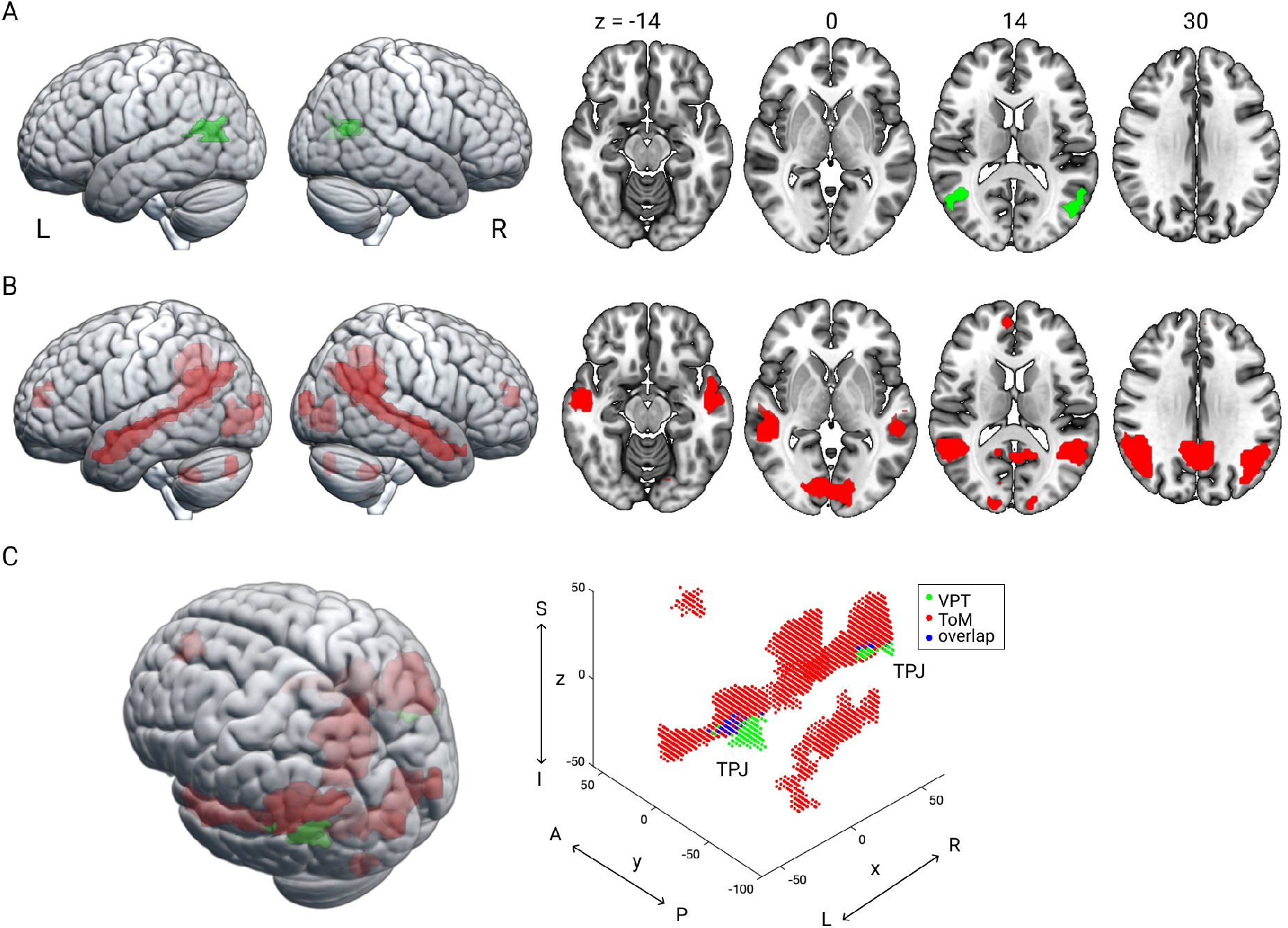
The activated regions on the surface and in horizontal planes displayed with MRIcroGL. (A) Regions with significantly higher activation during the VPT task than during the MR task. (B) Regions with significantly higher activation during the ToM task than during the control trials. (C) The activated voxels obtained during the VPT (3A in green, number of voxels = 3,124) and ToM (3B in red, number of voxels = 130) tasks were overlaid onto the 3D template brain of MRICroGL (left) and displayed with the 3D plot of MATLAB (right) in the same orientation. The blue voxels are the voxels overlapping between the VPT and ToM conditions (number of voxels = 69). The activation was considered significant for a threshold of p < .05 corrected for multiple comparisons for family-wise error (FWE) at the cluster-level with p < .001 uncorrected at the voxel level. L, left; R, right; S, superior; I, inferior; A, anterior; P, posterior. The activated areas are listed in Table 1. The unthresholded activation images in NIfTY format are available at https://doi.org/10.6084/m9.figshare.17046179

**Table 1:** Anatomical regions, peak voxel coordinates, and t-values of the observed activations.

| Anatomic region | voxels | MNI coordinates |  |  | *t*-value |
| --- | --- | --- | --- | --- | --- |
|  |  | x | y | z |  |
| *VPT > MR* |  |  |  |  |  |
| L TPJ | 130 | -51 | -67 | 14 | 5.52 |
| R TPJ | 69 | 54 | -64 | 14 | 4.78 |
| *ToM > control* |  |  |  |  |  |
| R middle temporal cortex | 1014 | 54 | 5 | -28 | 9.15 |
| R TPJ |  | 48 | -64 | 26 | 8.44 |
| L middle temporal cortex | 771 | -54 | -7 | -16 | 8.29 |
| L TPJ |  | -45 | -58 | 23 | 7.00 |
| M precuneus | 787 | 3 | -52 | 35 | 7.72 |
| R lingual gyrus | 443 | 6 | -82 | -4 | 7.23 |
| L lingual gyrus |  | -9 | -85 | -7 | 5.57 |
| L cerebellum | 45 | -21 | -79 | -34 | 6.02 |
| M anterior cingulate cortex | 59 | 6 | 53 | 11 | 5.58 |
| L mPFC |  | -9 | 47 | 23 | 5.33 |
| R mPFC |  | 6 | 56 | 20 | 4.76 |
| L cerebellum | 74 | -6 | -55 | -40 | 5.25 |
| R cerebellum |  | 9 | -46 | -40 | 4.80 |

MNI, Montreal Neurological Institute; VPT, visual perspective taking; MR, mental rotation; ToM, theory of mind; L, left hemisphere; R, right hemisphere; M, medial; TPJ, temporo-parietal junction; mPFC, medial prefrontal cortex.

To compare individual peaks within the TPJ, peak voxels were selected within the TPJ ROI mask. No significant activation (p < .05, uncorrected for the voxel level) was found within the TPJ in the left and right hemisphere of participants 3 and 2 during the VPT or ToM tasks. Therefore, these participants were excluded from the coordinate comparison. The peak coordinates were plotted using BrainNet Viewer (Xia et al., 2013). The peak voxels obtained during the ToM task were found in locations significantly more anterior (y-axis left hemisphere: t[27] = 3.06, p = .005, Cohen’s d = 0.59; y-axis right hemisphere: t[28] = 2.70, p = .012, Cohen’s d = 0.51) and dorsal (z-axis left hemisphere: t[27] = 3.52, p = .002, Cohen’s d = 0.68; z-axis right hemisphere: t[28] = 2.49, p = .019, Cohen’s d = 0.47) compared with the peak voxel positions in the bilateral TPJ for VPT tasks. There was no significant difference in the lateromedial direction, x-axis left hemisphere: t(27) = 0.85, p = .40, Cohen’s d = 0.16 and x-axis right hemisphere: t(28) = 0.53, p = .60, Cohen’s d = 0.10 (Fig. 4).

**Fig. 4:**
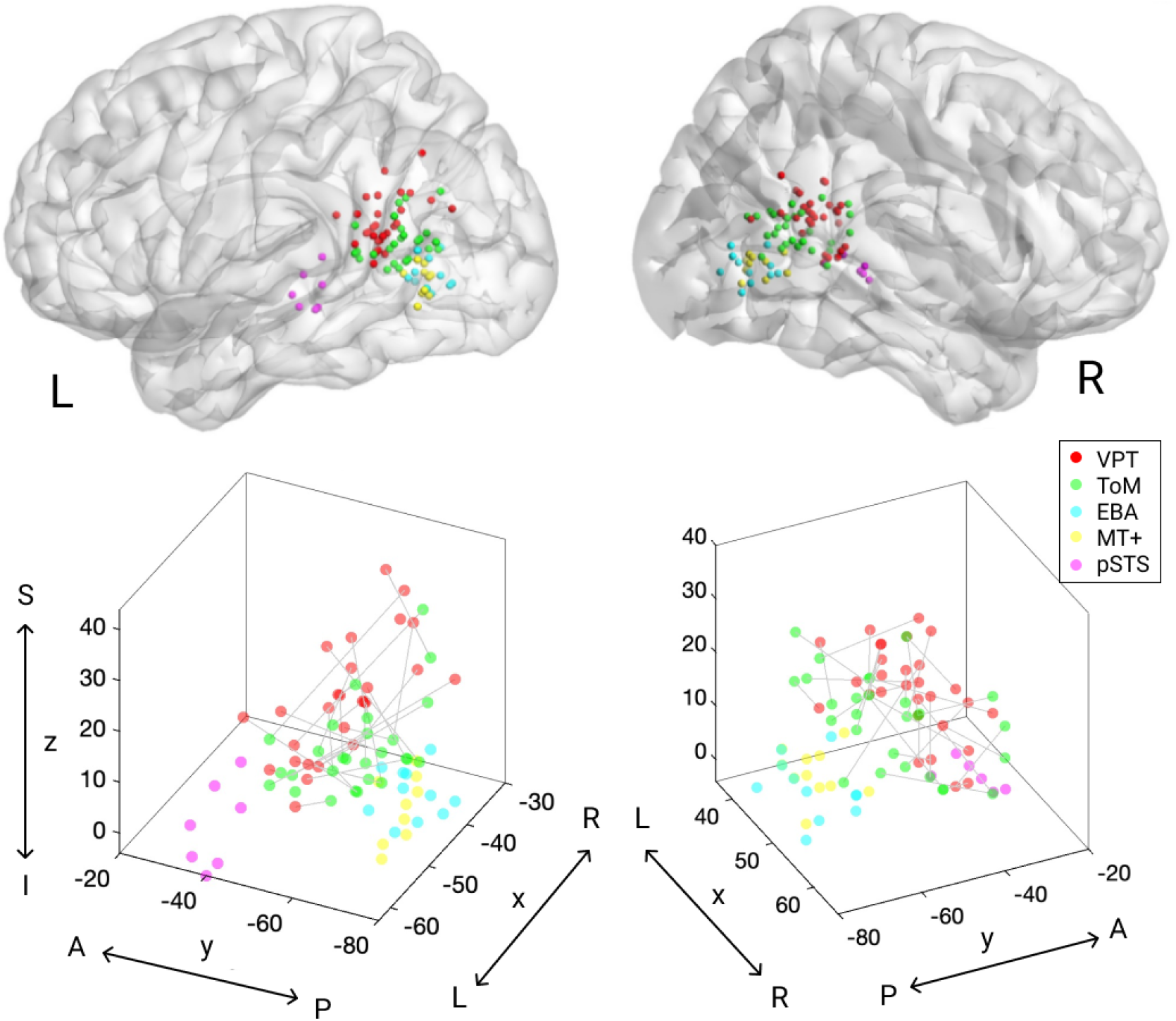
Individual peak voxels obtained during VPT (green) and ToM (red) task as well as for EBA (cyan), MT+ (yellow), and pSTS (magenta) displayed on the glass template brain using BrainNet Viewer (Xia et al. 2013) and the 3D plot on the MNI coordinates (lower panel). L, left; R, right; S, superior; I, inferior; A, anterior; P, posterior. The individual coordinate values are available at https://doi.org/10.6084/m9.figshare.17141867

We further used independent functional localizer scans to determine the peak localization in each participant for the EBA, MT+, and pSTS. All participants showed reliable EBA and MT+ activations in both the hemispheres, whereas we observed no activation in the pSTS from the left and right hemispheres for participants 5 and 6 and therefore did not include them in the analysis. We compared the individual peaks and found that these functional clusters were significantly spatially distinct from those obtained during VPT and ToM tasks in the x- and y-axes for both hemispheres (Fig. 4) (VPT vs. EBA y-axis: ts > 2.70, ps < 0.05, Cohen’s ds > 0.85; VPT vs. EBA z-axis: ts > 6.07, ps < 0.001, Cohen’s ds > 1.92; ToM vs. EBA y-axis: ts > 5.00, ps < 0.001, Cohen’s ds > 1.51; ToM vs. EBA z-axis: ts > 8.07, ps < 0.001, Cohen’s ds > 2.55; VPT vs. MT+ y-axis: ts > 2.32, ps < 0.05, Cohen’s ds > 0.73; VPT vs. MT+ z-axis: ts > 5.30, ps < 0.001, Cohen’s ds > 1.68; ToM vs. MT+ y-axis: ts > 6.31, ps < 0.001, Cohen’s ds > 1.90; ToM vs. MT+ z-axis: ts > 8.12, ps < 0.001, Cohen’s ds > 2.45; VPT vs. pSTS y-axis: ts > 7.48, ps < 0.005, Cohen’s ds > 3.74; VPT vs. pSTS z-axis: ts > 4.14, ps < 0.01, Cohen’s ds > 1.85; ToM vs. pSTS y-axis: ts > 2.62, ps < 0.06, Cohen’s ds > 1.31; ToM vs. pSTS z-axis: ts > 3.75, ps < 0.05, Cohen’s ds > 1.87).

## Discussion

This study investigated whether the same neural substrates in the TPJ are involved in VPT and ToM by directly comparing TPJ activation in individual participants. Behavioral data revealed that the response to the VPT task was significantly faster and more accurate than that to MR, whereas the response to the ToM task was slower than that to control tasks with no significant difference in task accuracy. The fMRI analyses revealed that the VPT and ToM tasks activated partially overlapped areas in the bilateral TPJ. By comparing the peak activations in each of the participants for both tasks, we found that peak voxels activated during the ToM task were located in a significantly more anterior and dorsal part of the TPJ than those activated by the VPT task. The activated areas were also spatially distinct from nearby functional modules (EBA, MT+, and pSTS), as shown by independent localizer scans. Our finding revealed neighboring but distinct representations of VPT and ToM within the TPJ.

We found focal activities in the bilateral TPJ induced by VPT when compared with MR-related activation. This TPJ activation was partially overlapped with the ToM activity, which corroborates the proposal that the VPT2 is a mentalizing task (Hamilton et al., 2009). Previous studies showed that the reorientation of the attention activates the TPJ (Corbetta et al., 2008). There have been controversies about whether the same TPJ areas are involved in the reallocation of attention and the ToM (Decety and Lamm, 2007; Mitchell, 2008; Young et al., 2010b; Geng and Vossel, 2013; Santiesteban et al., 2017). A meta-analysis revealed a substantial overlap between attentional reorientation and ToM in the right TPJ (rTPJ) (Decety and Lamm, 2007), whereas other studies showed that the posterior part of the rTPJ is involved in social cognition (Scholz et al., 2009; Krall et al., 2015). The TPJ activations observed in the present study are unlikely owing to the attentional processes, because VPT and MR tasks require bottom-up attention to visual cues and comparable mental spatial processing (Hamilton et al., 2009). The behavioral analyses showed faster and more accurate responses in the VPT task than in the MR task, indicating that the attentional demand or the global task difficulty was lower in the VPT task than in the MR task. In addition, the TPJ activation peak was closer to the coordinates obtained for ToM than that found in attentional reorientation (Decety and Lamm, 2007). For example, the coordinates [54 −64 14] in the right hemisphere (Table 1) are closer to the mean coordinates found in ToM [56, −54, 19] than those obtained in previous studies for attentional reorientation [55, −55, 26](Mitchell, 2008; Scholz et al., 2009). It should be noted that the visual cues were different between the VPT (avatar) and MR (edge color) tasks. Therefore, the observed activity might be a result of viewing the avatar. To overcome this problem, we conducted an EBA functional localizer, because EBA is involved in the visual perception of human bodies (Downing et al., 2001) and found that EBA activation was spatially distinct from the TPJ activation.

We not only found large areas activated by the ToM task, including the bilateral TPJ, but also the anterior part of the temporal cortex, the mPFC, the precuneus, and the lingual gyrus. Although previous studies indicated a dominance of the right hemisphere in ToM tasks (Sommer et al., 2007; Scholz et al., 2009; Young et al., 2010a), a bilateral involvement was also reported (Saxe and Kanwisher, 2003; Kobayashi et al., 2007), which is consistent with our result. The large area of activity in both hemispheres also agrees with previous neuroimaging studies on the ToM (Aichhorn et al., 2009; Lee and McCarthy, 2016; Ogawa et al., 2017), which showed activities in the mPFC, the precuneus, and the anterior part of the temporal cortex, in addition to the TPJ, constituting a core network for ToM performance (Frith and Frith, 1999; Amodio and Frith, 2006; Heyes and Frith, 2014; Schurz et al., 2014).

By comparing the whole-brain activations during the VPT and ToM tasks, we found partially overlapped voxels within the TPJ. This overlap was located within the angular gyrus and corresponded with the posterior part of the TPJ (TPJp), rather than the anterior TPJ (TPJa), which is consistent with the roles of the TPJp in social cognition, whereas the TPJa is involved in attentional reorientation (Decety and Lamm, 2007; Scholz et al., 2009; Mars et al., 2012; Carter and Huettel, 2013; Kubit and Jack, 2013). We further compared the peak activations of both functions in individual participants and found that the peak voxels induced by ToM were located significantly more anteriorly and dorsally within the bilateral TPJ compared with those observed during the VPT task. Both VPT and ToM share cognitive processes that require the mental representation of other agents, which is decoupled with the self-perspective of belief. Additionally, ToM necessitates a more complex representation of ones’ beliefs or thoughts for a longer time during narrative comprehension, whereas VPT requires a more visual and instantaneous representation of another’s perspectives. This functional discrepancy in representing ones’ minds might cause distinct peaks of activities during ToM and VPT tasks. Thus, it is not surprising that VPT-dependent activation is located ventrally to the early visual cortex for low-level visual processing, whereas the ToM-dependent activity is detected in a more anterior and ventral area with more complex cognitive components.

The VPT and ToM activity peaks were also found more anteriorly and dorsally than the activities in the EBA and pSTS, which are both involved in the social perception of others, the EBA of human body parts (Downing et al., 2001) and the pSTS of biological motion or detection of intentional agent (Chakrabarti and Baron-Cohen, 2006). These results suggest the existence of a functional gradient of social cognition, starting from the low-level detection of other agents in the pSTS and EBA and ending in the more anterior and dorsal areas activated by ToM and VPT with the high-level mental representation of the other agent decoupled from oneself. Our proposal is in line with the nexus model of the TPJ, which proposes that the basic social perception begins in the lateral occipital cortex (EBA and pSTS) and then converges dorsally into the abstract representation for social cognition (Carter and Huettel, 2013), together with functional heterogeneity in the TPJ (Lee and McCarthy, 2016). The TPJ is also associated with other types of perspective taking of others under complex social interactions, including joint action (Abe et al., 2019), charitable giving (Tusche et al., 2016), strategic competitive interaction (Ogawa and Kameda, 2020), social judgment or decision making (Young et al., 2010a; Kameda et al., 2016). Further studies are needed to understand the relationships between these functions and clarify the neural mechanisms and functional gradients within the TPJ.

In summary, the present study investigated whether VPT and ToM have the same neural substrates in the TPJ by directly comparing the activation patterns of individual participants using fMRI with a within-subjects design. Our findings revealed that VPT and ToM have neighboring but distinct representations, indicating the functional heterogeneity of social cognition within the TPJ.

## Acknowledgments

This work was supported by JSPS KAKENHI Grant Numbers 17H01016 & 19H00634 to K.O. The authors would like to thank Enago (www.enago.jp) for the English language review.

## Author contributions

K.O. and Y.M. designed research, performed research, and analyzed data; K.O. wrote the paper.

## Code accessibility

The computer code is available from the corresponding author upon reasonable request.

## References

Abe MO, Koike T, Okazaki S, Sugawara SK, Takahashi K, Watanabe K, Sadato N (2019) Neural correlates of online cooperation during joint force production. Neuroimage 191:150–161.

Agarwal SM, Shivakumar V, Kalmady SV, Danivas V, Amaresha AC, Bose A, Narayanaswamy JC, Amorim MA, Venkatasubramanian G (2017) Neural correlates of a perspective-taking task using in a realistic three-dimmensional environment based task: A pilot functional magnetic resonance imaging study. Clin Psychopharmacol Neurosci 15:276–280.

Aichhorn M, Perner J, Kronbichler M, Staffen W, Ladurner G (2006) Do visual perspective tasks need theory of mind? Neuroimage 30:1059–1068.

Aichhorn M, Perner J, Weiss B, Kronbichler M, Staffen W, Ladurner G (2009) Temporo-parietal junction activity in theory-of-mind tasks: Falseness, beliefs, or attention. J Cogn Neurosci 21:1179–1192.

Alcalá-López D, Smallwood J, Jefferies E, Van Overwalle F, Vogeley K, Mars RB, Turetsky BI, Laird AR, Fox PT, Eickhoff SB, Bzdok D (2018) Computing the Social Brain Connectome Across Systems and States. Cereb Cortex 28:2207–2232.

Amodio DM, Frith CD (2006) Meeting of minds: the medial frontal cortex and social cognition. Nat Rev Neurosci 7:268–277.

Apperly IA, Samson D, Chiavarino C, Humphreys GW (2004) Frontal and temporo-parietal lobe contributions to theory of mind: neuropsychological evidence from a false-belief task with reduced language and executive demands. J Cogn Neurosci 16:1773–1784.

Arora A, Schurz M, Perner J (2017) Systematic Comparison of Brain Imaging Meta-Analyses of ToM with vPT. Biomed Res Int 2017:1–12.

Bzdok D, Langner R, Schilbach L, Jakobs O, Roski C, Caspers S, Laird AR, Fox PT, Zilles K, Eickhoff SB (2013) Characterization of the temporo-parietal junction by combining data-driven parcellation, complementary connectivity analyses, and functional decoding. Neuroimage 81:381–392.

Carter RM, Huettel SA (2013) A nexus model of the temporal – parietal junction. Trends Cogn Sci 17:328–336.

Chakrabarti B, Baron-Cohen S (2006) Empathizing: neurocognitive developmental mechanisms and individual differences. Prog Brain Res 156:403–417.

Corbetta M, Patel G, Shulman GL (2008) The reorienting system of the human brain: from environment to theory of mind. Neuron 58:306–324.

Corradi-Dell’Acqua C, Ueno K, Ogawa A, Cheng K, Rumiati RI, Iriki A (2008) Effects of shifting perspective of the self: An fMRI study. Neuroimage 40:1902–1911.

Decety J, Lamm C (2007) The role of the right temporoparietal junction in social interaction: How low-level computational processes contribute to meta-cognition. Neuroscientist 13:580–593.

Dodell-Feder D, Koster-Hale J, Bedny M, Saxe R (2011) fMRI item analysis in a theory of mind task. Neuroimage 55:705–712.

Downing PE, Jiang Y, Shuman M, Kanwisher N (2001) A cortical area selective for visual processing of the human body. Science 293:2470–2473.

Flavell JH (1977) The development of knowledge about visual perception. Nebr Symp Motiv 25:43–76.

Flavell JH, Everett BA, Croft K, Flavell ER (1981) Young children’s knowledge about visual perception: Further evidence for the Level 1–Level 2 distinction. Dev Psychol 17:99–103.

Frith CD, Frith U (1999) Interacting minds--a biological basis. Science 286:1692–1695.

Geng JJ, Vossel S (2013) Re-evaluating the role of TPJ in attentional control: contextual updating? Neurosci Biobehav Rev 37:2608–2620.

Grossman E, Donnelly M, Price R, Pickens D, Morgan V, Neighbor G, Blake R (2000) Brain areas involved in perception of biological motion. J Cogn Neurosci 12:711–720.

Gunia A, Moraresku S, Vlček K (2021) Brain mechanisms of visuospatial perspective-taking in relation to object mental rotation and the theory of mind. Behav Brain Res 407:113247.

Gzesh SM, Surber CF (1985) Visual perspective-taking skills in children. Child Dev 56:1204–1213.

Hamilton AF de C, Brindley R, Frith U (2009) Visual perspective taking impairment in children with autistic spectrum disorder. Cognition 113:37–44.

Hatta T, Nakatsuka Z (1975) Handedness inventory. In: Papers on Celebrating 63rd Birthday of Prof.Ohnishi, pp 224–245. Osaka City University.

Heyes CM, Frith CD (2014) The cultural evolution of mind reading. Science 344:1243091.

Hirai M, Muramatsu Y, Nakamura M (2018) Role of the Embodied Cognition Process in Perspective-Taking Ability During Childhood. Child Dev Available at: http://doi.wiley.com/10.1111/cdev.13172.

Hong Y-W, Yoo Y, Han J, Wager TD, Woo C-W (2019) False-positive neuroimaging: Undisclosed flexibility in testing spatial hypotheses allows presenting anything as a replicated finding. Neuroimage 195:384–395.

Kameda T, Inukai K, Higuchi S, Ogawa A, Kim H, Matsuda T, Sakagami M (2016) Rawlsian maximin rule operates as a common cognitive anchor in distributive justice and risky decisions. Proc Natl Acad Sci U S A 113:11817–11822.

Kanske P, Böckler A, Trautwein FM, Singer T (2015) Dissecting the social brain: Introducing the EmpaToM to reveal distinct neural networks and brain-behavior relations for empathy and Theory of Mind. Neuroimage 122:6–19.

Kessler K, Rutherford H (2010) The two forms of visuo-spatial perspective taking are differently embodied and subserve different spatial prepositions. Front Psychol 1:1–12.

Kobayashi C, Glover GH, Temple E (2007) Children’s and adults’ neural bases of verbal and nonverbal “theory of mind.” Neuropsychologia 45:1522–1532.

Krall SC, Rottschy C, Oberwelland E, Bzdok D, Fox PT, Eickhoff SB, Fink GR, Konrad K (2015) The role of the right temporoparietal junction in attention and social interaction as revealed by ALE meta-analysis. Brain Struct Funct 220:587–604.

Kubit B, Jack AI (2013) Rethinking the role of the rTPJ in attention and social cognition in light of the opposing domains hypothesis: findings from an ALE-based meta-analysis and resting-state functional connectivity. Front Hum Neurosci 7:323.

Lee SM, McCarthy G (2016) Functional Heterogeneity and Convergence in the Right Temporoparietal Junction. Cereb Cortex 26:1108–1116.

Mars RB, Sallet J, Schüffelgen U, Jbabdi S, Toni I, Rushworth MFS (2012) Connectivity-based subdivisions of the human right “temporoparietal junction area”: evidence for different areas participating in different cortical networks. Cereb Cortex 22:1894–1903.

Martin AK, Kessler K, Cooke S, Huang J, Meinzer M (2020) The Right Temporoparietal Junction Is Causally Associated with Embodied Perspective-taking. Journal of Neuroscience 40:3089–3095.

Michelon P, Zacks JM (2006) Two kinds of visual perspective taking. Percept Psychophys 68:327–337.

Mitchell JP (2008) Activity in right temporo-parietal junction is not selective for theory-of-mind. Cereb Cortex 18:262–271.

Ogawa A, Kameda T (2020) Dissociable roles of left and right temporoparietal junction in strategic competitive interaction. Soc Cogn Affect Neurosci 14:1037–1048.

Ogawa A, Yokoyama R, Kameda T (2017) Development of a Japanese version of a theory-of-mind functional localizer for functional magnetic resonance imaging. The Japanese Journal of Psychology 88:366–375.

Okamoto Y, Kosaka H, Kitada R, Seki A, Tanabe HC, Hayashi MJ, Kochiyama T, Saito DN, Yanaka HT, Munesue T, Ishitobi M, Omori M, Wada Y, Okazawa H, Koeda T, Sadato N (2017) Age-dependent atypicalities in body- and face-sensitive activation of the EBA and FFA in individuals with ASD. Neurosci Res 119:38–52.

Oldfield RC (1971) The assessment and analysis of handedness: the Edinburgh inventory. Neuropsychologia 9:97–113.

Otsuka Y, Osaka N, Ikeda T, Osaka M (2009) Individual differences in the theory of mind and superior temporal sulcus. Neurosci Lett 463:150–153.

Pearson A, Ropar D, de C. Hamilton AF (2013) A review of visual perspective taking in autism spectrum disorder. Front Hum Neurosci 7:1–10.

Peuskens H, Vanrie J, Verfaillie K, Orban GA (2005) Specificity of regions processing biological motion. Eur J Neurosci 21:2864–2875.

Samson D, Apperly IA, Chiavarino C, Humphreys GW (2004) Left temporoparietal junction is necessary for representing someone else’s belief. Nat Neurosci 7:499–500.

Samson D, Apperly IA, Kathirgamanathan U, Humphreys GW (2005) Seeing it my way: a case of a selective deficit in inhibiting self-perspective. Brain 128:1102–1111.

Santiesteban I, Banissy MJ, Catmur C, Bird G (2015) Functional lateralization of temporoparietal junction - imitation inhibition, visual perspective-taking and theory of mind. Eur J Neurosci 42:2527–2533.

Santiesteban I, Kaur S, Bird G, Catmur C (2017) Attentional processes, not implicit mentalizing, mediate performance in a perspective-taking task: Evidence from stimulation of the temporoparietal junction. Neuroimage 155:305–311.

Saxe R, Kanwisher N (2003) People thinking about thinking peopleThe role of the temporo-parietal junction in “theory of mind.” Neuroimage 19:1835–1842.

Scholz J, Triantafyllou C, Whitfield-Gabrieli S, Brown EN, Saxe R (2009) Distinct regions of right temporo-parietal junction are selective for theory of mind and exogenous attention Lauwereyns J, ed. PLoS One 4:e4869.

Schurz M, Aichhorn M, Martin A, Perner J (2013) Common brain areas engaged in false belief reasoning and visual perspective taking: a meta-analysis of functional brain imaging studies. Front Hum Neurosci 7:712.

Schurz M, Kronbichler M, Weissengruber S, Surtees A, Samson D, Perner J (2015) Clarifying the role of theory of mind areas during visual perspective taking: Issues of spontaneity and domain-specificity. Neuroimage 117:386–396.

Schurz M, Radua J, Aichhorn M, Richlan F, Perner J (2014) Fractionating theory of mind: a meta-analysis of functional brain imaging studies. Neurosci Biobehav Rev 42:9–34.

Shepard RN, Metzler J (1971) Mental rotation of three-dimensional objects. Science 171:701–703.

Sommer M, Döhnel K, Sodian B, Meinhardt J, Thoermer C, Hajak G (2007) Neural correlates of true and false belief reasoning. Neuroimage 35:1378–1384.

Spiridon M, Fischl B, Kanwisher N (2006) Location and spatial profile of category-specific regions inhuman extrastriate cortex. Hum Brain Mapp 27:77–89.

Tootell RB, Reppas JB, Kwong KK, Malach R, Born RT, Brady TJ, Rosen BR, Belliveau JW (1995) Functional analysis of human MT and related visual cortical areas using magnetic resonance imaging. J Neurosci 15:3215–3230.

Tusche A, Böckler A, Kanske P, Trautwein FM, Singer T (2016) Decoding the charitable brain: Empathy, perspective taking, and attention shifts differentially predict altruistic giving. Journal of Neuroscience 36:4719–4732.

Wilms M, Eickhoff SB, Specht K, Amunts K, Shah NJ, Malikovic A, Fink GR (2005) Human V5/MT+: Comparison of functional and cytoarchitectonic data. Anat Embryol 210:485–495.

Xia M, Wang J, He Y (2013) BrainNet Viewer: a network visualization tool for human brain connectomics Csermely P, ed. PLoS One 8:e68910.

Young L, Camprodon JA, Hauser M, Pascual-Leone A, Saxe R (2010a) Disruption of the right temporoparietal junction with transcranial magnetic stimulation reduces the role of beliefs in moral judgments. Proc Natl Acad Sci U S A 107:6753–6758.

Young L, Dodell-Feder D, Saxe R (2010b) What gets the attention of the temporo-parietal junction?An fMRI investigation of attention and theory of mind. Neuropsychologia 48:2658–2664.

